# Multielectrode Cortical Stimulation Selectively Induces Unidirectional Wave Propagation

**DOI:** 10.1101/2020.11.28.402289

**Authors:** Alma Halgren, Zarek Siegel, Ryan Golden, Maxim Bazhenov

## Abstract

Cortical stimulation is emerging as an experimental tool in basic research and a promising therapy for a range of neuropsychiatric conditions. As multielectrode arrays enter clinical practice, the possibility of using spatiotemporal patterns of electrical stimulation to induce desired physiological patterns has become theoretically possible, but in practice can only be implemented by trial-and-error because of a lack of predictive models. Experimental evidence increasingly establishes travelling waves as fundamental to cortical information-processing, but we lack understanding how to control wave properties despite rapidly improving technologies. This study uses a hybrid biophysical-anatomical and neural-computational model to predict and understand how a simple pattern of cortical surface stimulation could induce directional traveling waves via asymmetric activation of inhibitory interneurons. It reveals local circuit mechanisms to control spatiotemporal cortical dynamics and predicts interventions that can be developed to treat a broad range of cognitive disorders.

## INTRODUCTION

Brain stimulation is widely used in both experimental and clinical settings. In basic research it is used to probe neural function by disrupting or hyperactivating local brain processing^1–3^. In clinical settings, direct manipulation of activity via stimulation has also been shown to be an effective in treatment of several neural and psychiatric disorders. Deep brain stimulation (DBS) has been successful in the treatment of movement disorders like Parkinson’s disease^4–6^, depression^7,8^ and obsessive-compulsive disorder^9,10^.Superficial cortical stimulation is an effective therapy for epileptic^11^ and stroke patients^12^. Increasingly, electrical stimulation has also shown promise in both the restoration and enhancement of critical cognitive functions such as memory. DBS has been shown experimentally to enhance memory encoding when applied during learning^13^ and closed-loop stimulation protocols have proven to be effective during periods of poor memory encoding as well as during memory recall^14–16^.

While brain stimulation is sometimes conceptualized as disrupting pathological activity in order to restore normal activity, increasingly the explicit goal is to directly generate normal activity. Experimental evidence supports travelling waves as critical to normal brain activity. These propagating waves are fundamental to brain information-processing as they coordinate neural behavior across all spatial scales, from within-layer to whole-brain interactions, as well as across temporal scales, from tens to hundreds of milliseconds. By mediating communication across multiple brain areas, propagating activity putatively performs a variety of cognitive functions such as long-term memory consolidation and the processing of visual stimuli^17^. For example, sleep spindles travelling across the cortical surface at multiple scales have been hypothesized to synchronize convergent co-firing, resulting in spike-timing dependent plasticity and consequent memory consolidation^19^. Similarly, the alpha rhythm which modulates visual processing appears to be a travelling wave from association to primary areas^18^. Accordingly, the ability to predict and control traveling waves has far-reaching implications for improving and controlling cognitive function.

Currently, there is no method of reliably generating directional traveling waves with electrical stimulation. Past efforts to develop new paradigms of stimulation which reinstate particular brain activity states have largely depended upon trial and error. Recently, we described a method for modeling the effects of cortical stimulation that enables one to predict the consequences of stimulation in silico, and thereby develop stimulation protocols that achieve desired results in vivo^20^. This earlier study was limited to the effects of stimulation of a single electrode and thus did not evoke directional propagation. In this new study, we describe an initial attempt to model a multielectrode stimulation paradigm that produces unidirectional traveling waves in the cortex. With multielectrode arrays increasingly entering clinical practice^21^, our model harnesses the additional complexity and control during stimulation that multielectrode protocols allow for.

The modeling approach includes two phases. In the first phase, biophysical model is used to predict spiking probability in response to a spatially-varied electric field potential in reconstructed rat somatosensory cortical neurons obtained from NeuroMorpho.org^22^. We found that the hyperpolarization or depolarization of individual neurons varied according to cell type and cortical depth, and also varied with respect to the polarity of the applied electric field. The diversity in excitation response underlies the propagating wave activity that we observed in the second phase of the model. In the second phase, we constructed a Hodgkin-Huxley model of cortical network composed of multiple interconnected cortical columns, each containing a circuit of inhibitory and excitatory cells across cortical layers. Approximating stimulation effects using the activation probabilities calculated in phase one, we found that fast inhibitory activity, coupled with excitatory cells’ preference for cathodal stimulation, resulted in a unidirectionally propagating wave of activity. These results develop the previously published model for application to a new and potentially clinically-relevant domain, suggesting a simple multi-electrode pattern for evoking traveling waves and providing testable predictions for experimental confirmation and parameter optimization.

## RESULTS

Building on existing *in silico* models that simulated the effects of cortical surface stimulation from a single electrode^23^, we sought to explore the spatial dynamics of stimulation by modeling an asymmetrical three-electrode configuration (Fig 1a). The paper is organized as follows. We first calculated electrical field created by the system of three electrodes. Next, we estimated the activation probability for each cell type and cortical layer by computing the activating function in biophysical reconstructions of axonal arbors. We then constructed a cortical microcircuit model with Hodgkin-Huxley dynamics to model the network effects of stimulation based on the previously calculated spiking probabilities.

**Figure 1.**
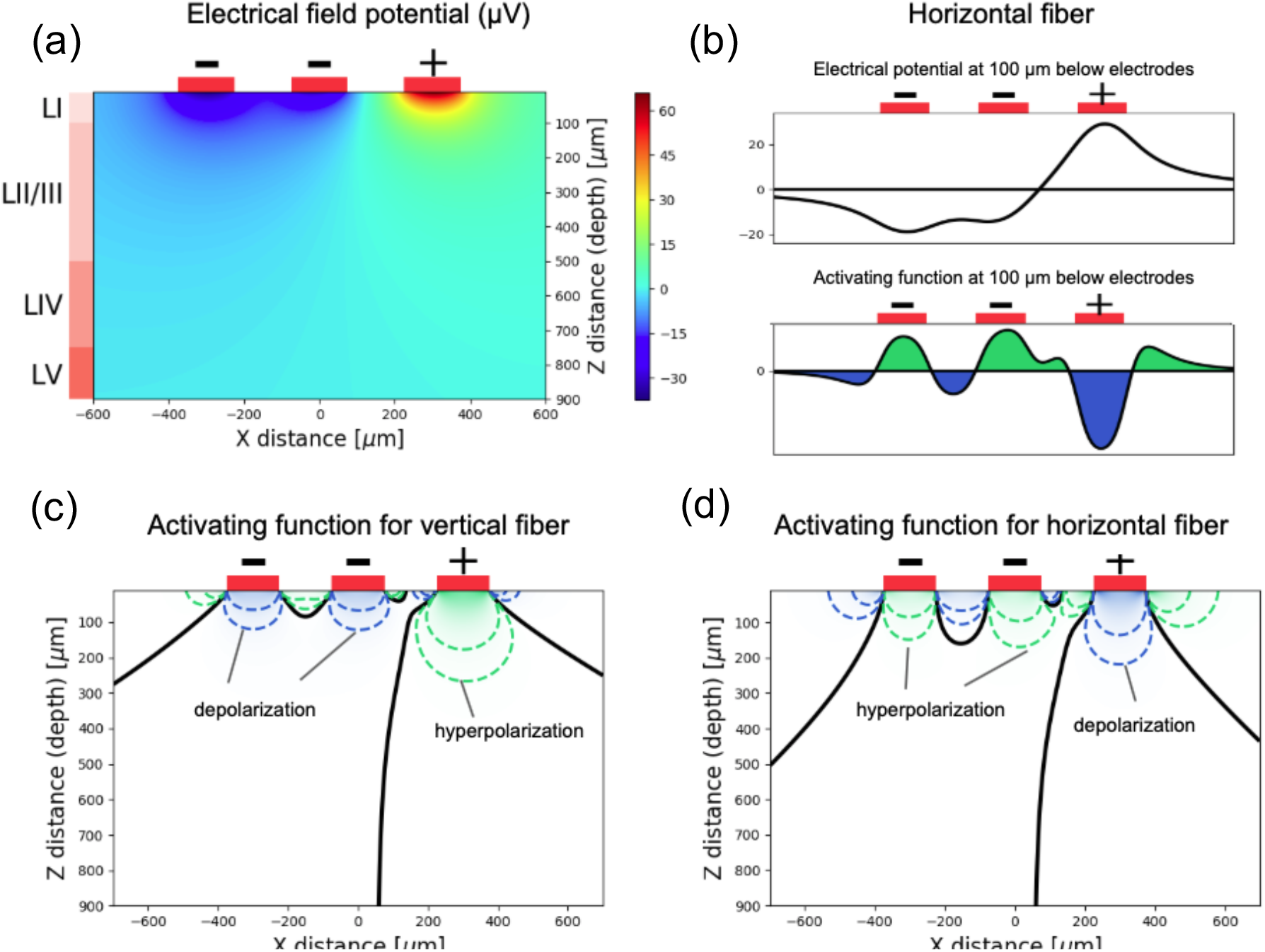
Electric potential and activating function in the plane Y=0. 1a. Schematic of electrode configuration and electric potential in the plane Y=0. Each electrode is square-shaped and measures 150 μm by 150 μm, and the spacing between each electrode is also 150 μm. The two leftmost electrodes are negative and deliver negative 75 μA of current each, while the rightmost electrode is positive and delivers 150 μA of current, and the stimulation period is 200 μs. Layer depths are approximated from experimental measurements of rat cortex^24,25,71,84^ as follows: layer I is 0 to 100 μm, layer II/III is 100 to 500 μm, layer IV is 500 to 750 μm, and layer V is 750 to 900 μm. 1b. Axial potential along horizontal axonal fiber located 100 um below the cortical surface. The axon is depolarized when the activating function is below zero and hyperpolarized when the activating function is above zero. 1c and 1d. As a result of 1b, horizontal fibers are activated by the cathode and hyperpolarized by the anodes (1d). The opposite is true for vertical fibers (1c). The vast majority of direct stimulation occurs in layers I and II/III due to the decay of electric potential with depth.

### Cell activation results from a combination of morphology (cell type) and depth within the column

The applied electric field potential generated by the system of three electrodes (assuming homogenous tissue) is shown in Fig 1a. To estimate the probability of specific cell types being activated by stimulation, we simulated the various cell types based on three-dimensional (3D) morphological reconstructions of neurons derived from electron microscopy available from NeuroMorpho.org^22^. The excitatory cells we considered were pyramidal cells across layers II-V, while the inhibitory neurons included basket cells and Martinotti cells across layers II-V in addition to layer I interneurons (Table S1). Example reconstructions as well as average axon density plots per cell type/cortical layer (Fig 2) demonstrate the significant differences in axonal arborization and density among the different cell types, as well as between cells of the same type based in different layers.

**Figure 2.**
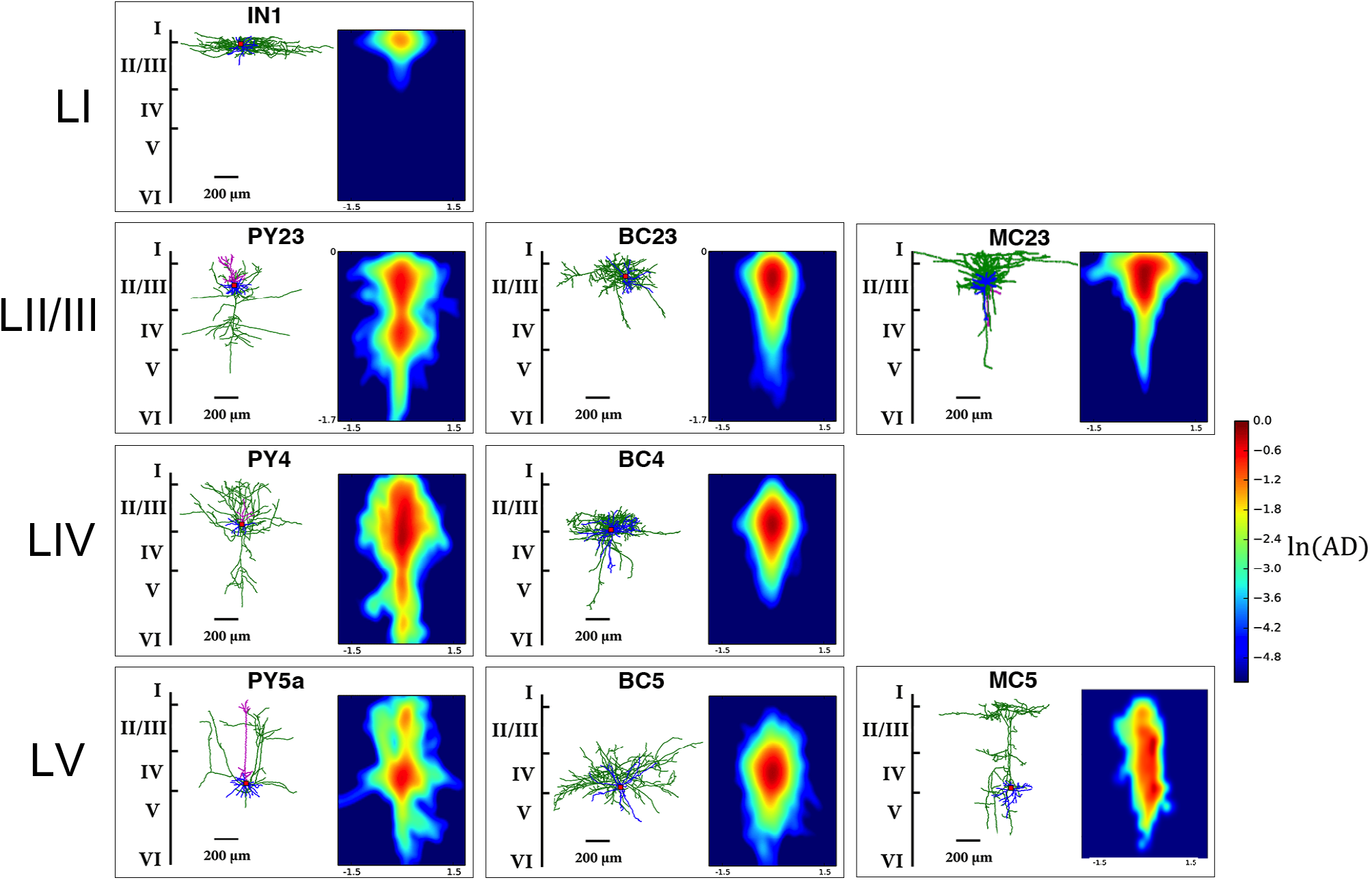
Representative reconstructions and averaged axon density for neuronal cell types modeled. For each neuronal cell type employed in the cortical microcircuit model, we plot both a single representative anatomical reconstruction as well as an averaged axonal density heatmap for all the reconstructions of that type. Cells are arranged by layer and type, where the first row depicts layer I inhibitory neurons. The representative LI interneuron is a horizontal cell, but we average across small, descending, and horizontal LI interneurons to obtain the axonal heatmap as well as in our analyses. The following rows depict pyramidal, basket, and Martinotti cells across layers II/III, IV, and V (LVa for pyramidal cells). In axonal density (AD) plots, color indicates the averaged density computed using all available reconstructions for the given cell type. The color scale is logarithmic for visual clarity. AD gives a sense of the general orientation and density of axon branches for each cell type, which is key to understanding subsequently computed activation probabilities.

The hyperpolarization or depolarization of a neuronal fiber within a constant electric field can be modeled with one-dimensional cable theory in conjunction with the activating function. The activating function (see Methods) computes the net transmembrane current generated by external stimulation (while ignoring pre-existing synaptic currents). According to one-dimensional cable theory, the activating function is the second-order spatial derivative of the electric potential along the neuronal fiber. The case of a perfectly horizontal fiber is shown in Figure 1b. Through this, we can draw relationships between the orientation and excitation of a fiber in response to a given stimulation polarity. Indeed, horizontal fibers were depolarized by cathodal stimulation and hyperpolarized by anodal stimulation, while in contrast vertical fibers were hyperpolarized by cathodal stimulation and depolarized by anodal stimulation (Figures 1d and 1c, respectively). While each neuron has unique axonal fibers that span three-dimensional space, these maps of activation and suppression zones for orthogonal axonal orientations give us insight into how each cell type will behave across the stimulated space given its average axon density and orientation (Fig 2).

We next calculated the spiking probability in response to the applied electric field potential for each cell type/cortical layer by averaging across the activating function results of their respective cell reconstructions; each cell reconstruction was shuffled by rotating and shifting along the vertical axis and multiple reconstructions were considered for each cell type (see Methods and Komarov, et al.^20^ for details). This calculation compares the overall excitability of each reconstruction to an experimentally-derived threshold (*f*_*th*_ = 3 pA/μm^2^) to determine probability of spiking. This threshold was set twenty times higher for unmyelinated cell types (Martinotti cells and layer I interneurons) compared to myelinated cell types (pyramidal and basket cells) since unmyelinated fibers are relatively unexcitable and lack nodes of Ranvier^24–26^.

The results of these calculations are shown in Figure 3. The average axonal density heatmaps in Figure 2 as well as the presence or absence of myelination explain the variation of activation responses across cell types and cortical layers.

**Figure 3.**
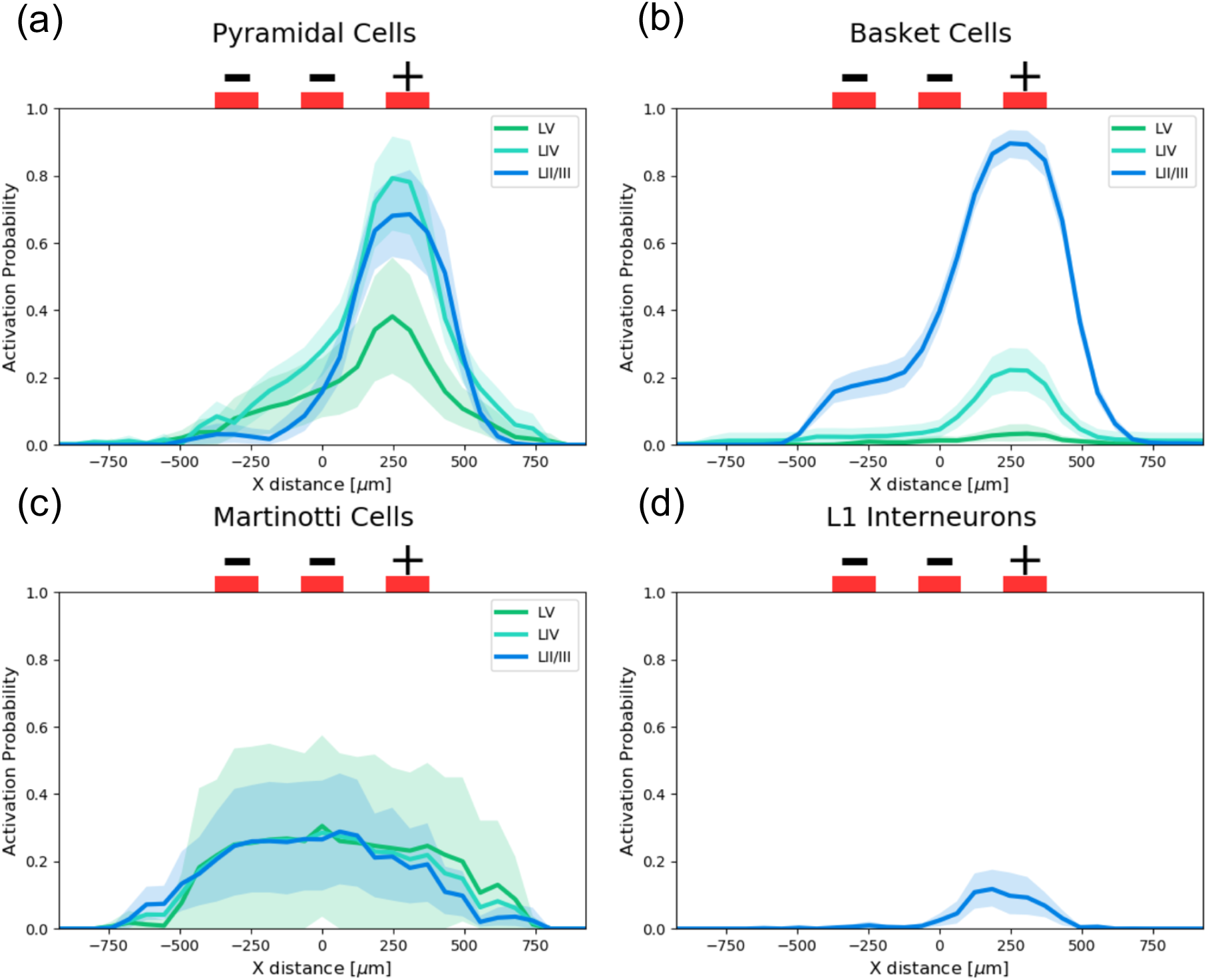
Probability of spiking as a function of horizontal distance from the center electrode for each cell type and cortical layer. Average (solid line) cell spiking probability and 95% confidence intervals (shaded region) for each reconstruction was calculated for soma locations across the entire X-Z plane of its cortical layer by averaging spiking probability across rotations and vertical shifts of all cell reconstructions. Activation probabilities were calculated with a cathodal stimulation current of 150 μA and an anodal stimulation current of −75 μA per electrode over a 200 μs stimulation period. Pyramidal cells (excitatory) and Basket cells (inhibitory) are highly activated by cathodal stimulation and are minimally activated by anodal stimulation due to their myelination and horizontally oriented axonal arbors. Martinotti cells (inhibitory) are activated by all electrodes with a slight preference for the anodes but lack myelination and thus show less activation overall. LI interneurons are also unmyelinated and are minimally excited by stimulation.

Across all layers, pyramidal cells were strongly activated by the cathode and minimally activated by the anodes but showed decreasing intensity of stimulation response with increasing depth (Fig 3a). As shown in Figure 2, all pyramidal cells vertically span the cortical layers regardless of soma position. However, Layer II/III and LIV pyramidal cells exhibit significant horizontal axonal density close to the cortical surface and thus responded more strongly to stimulation overall (and to cathodal stimulation in particular), whereas the bulk of Layer Va pyramidal axons lie in deeper layers and lack the superficial axonal density to be adequately stimulated above threshold. Strong overall pyramidal response to stimulation is due to their myelinate axons.

Basket cells also exhibited a strong preference for cathodal stimulation and little activation underneath the anodes (Fig 3b). However, their responses were significantly more tiered according to cortical layer as compared to pyramidal cells because basket cell arborization is localized within the same layer as the soma (Figure 2). Their preference for cathodal stimulation is due to their largely horizontal axonal arbor that stretches out within each layer. Basket cells were the only myelinated inhibitory cell type in our model and therefore demonstrated a significantly stronger spiking response overall relative to Martinotti or layer I interneurons.

Martinotti cells across all cortical layers are moderately activated by both cathodal and anodal stimulation but showed a slight preference for the latter (Fig 3c). This is because all Martinotti cells make universal connections with pyramidal cells via layer I (Fig 2), and therefore the majority of their arborizations lie in vertical axonal fibers connecting the soma to layer I, with additional density spread out horizontally across layer I. However, they exhibited a dampened stimulation response overall due to their unmyelinated axons.

Lastly, since layer I axon fibers are unmyelinated and stay localized to layer I (resulting in mainly horizontal arborization), layer I interneurons to show a slight preference for cathodal stimulation but little activation overall (Fig 3d).

### Cortical microcircuit model shows directional propagation when stimulated with three electrode array

In the previous section, we estimated the activation probabilities of isolated neurons within an applied electric field. To understand how stimulation affects the dynamics between neurons and ultimately the overall dynamics of the cortex, we constructed and stimulated a network model of the cortex using simplified neuron models and previously-calculated activation probabilities. Each cortical column was modeled as a canonical microcircuit containing the same cell types and cortical layers as the biophysical analysis above; cross-columnar connections were formed only between excitatory cells within the same layer. A schematic of the cell types and synaptic connections is shown in Figure 4.

**Figure 4.**
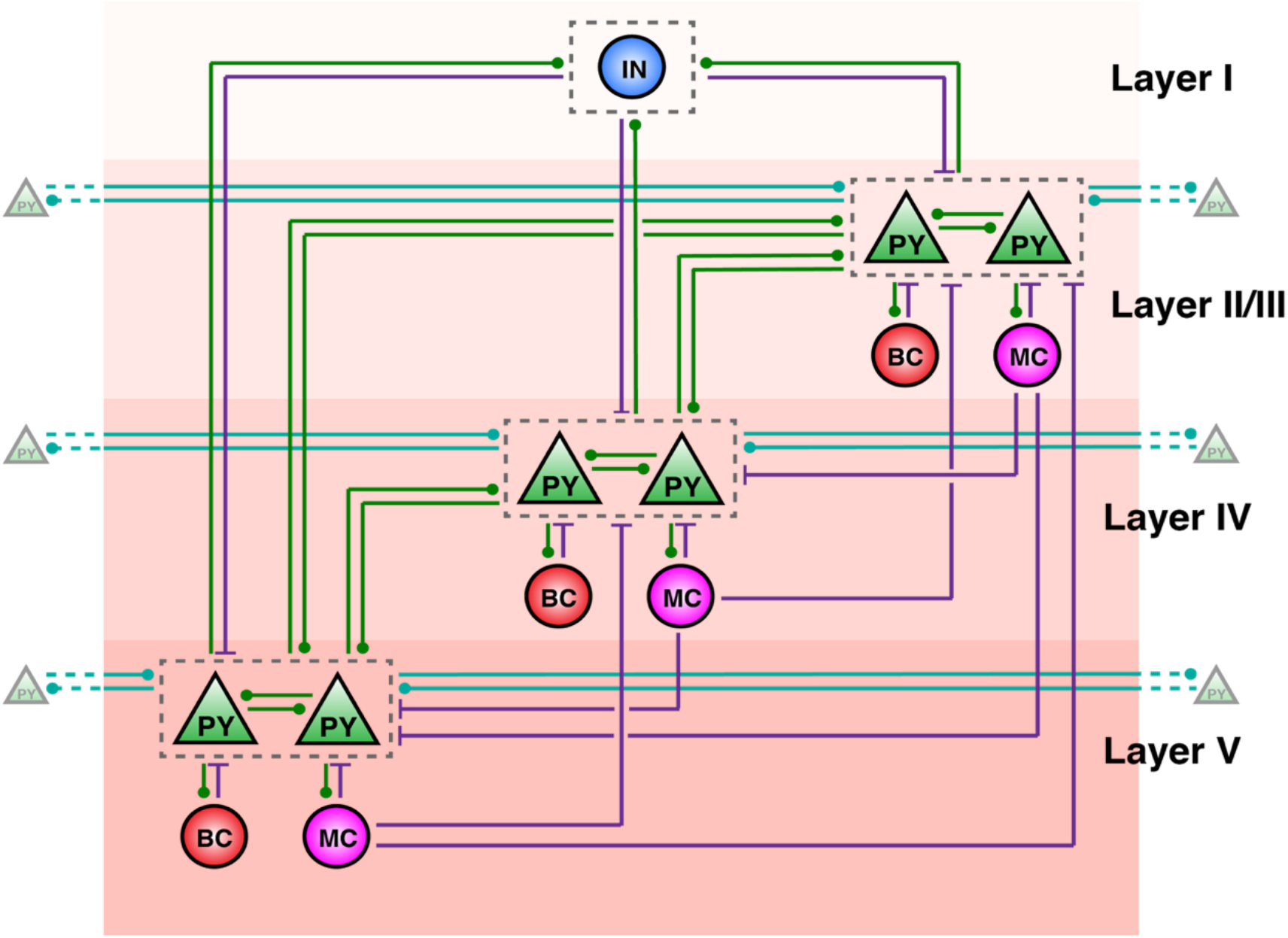
Microcircuit diagram of a single cortical column in modeled network. This depiction of a single cortical column details the cell types/cortical layers and the synaptic connections between them in our microcircuit network model. Circular labels (IN – LI interneurons; BC – basket cells; MC – Martinotti cells) indicate inhibitory neurons, while triangular labels (PY – pyramidal cells) indicate excitatory neurons. Green connections indicate excitatory synapses, with the circular end indicating the postsynaptic cell and the unlabeled end indicating the presynaptic cell. Purple connections indicate inhibitory synapses, with the perpendicular line indicating the postsynaptic end. Teal-colored connections are also excitatory but represent connections from pyramidal cells to others in adjacent columns (cross-columnar synapses). The only cross-column synapses present are within-layer pyramidal-pyramidal excitatory connections to adjacent columns. Layer I includes only inhibitory interneurons, which inhibit pyramidal cells in all three deeper layers and are reciprocally excited by the same cells. Each of the deeper layers contains pyramidal, basket, and Martinotti cells. Within a cortical column, pyramidal cells reciprocally excite other pyramidal cells in their same layer as well as across cortical layers. Basket cells act as local interneurons as they only inhibit and are excited by pyramidal cells within their own layer. In contrast, while Martinotti cells are only excited by pyramidal cells within their own layer, they universally inhibit pyramidal cells across all layers. In this model, inhibitory cells receive only excitatory synaptic inputs. The number of neurons in each column is listed in Table S2.

Following a brief stimulation period in which the activation probabilities from the biophysical calculations were applied to the model, the network was allowed to run without any external input for 500 ms, behaving according to synaptic interactions between neurons. A raster plot and voltage and conductance traces of one microcircuit trial are shown in Figures 5 and 6. The trial shown is one example of the general behavior of the microcircuit in the majority of trials (Fig 5c) in which spiking activity, particularly from LII/III pyramidal cells, propagates to the rightward columns but not past the leftmost electrode. Given this unique spiking activity and the biological importance of LII/III pyramidal cells in mediating communication across cortical regions, we chose to focus our analyses on the network behavior of LII/III pyramidal cells.

**Figure 5.**
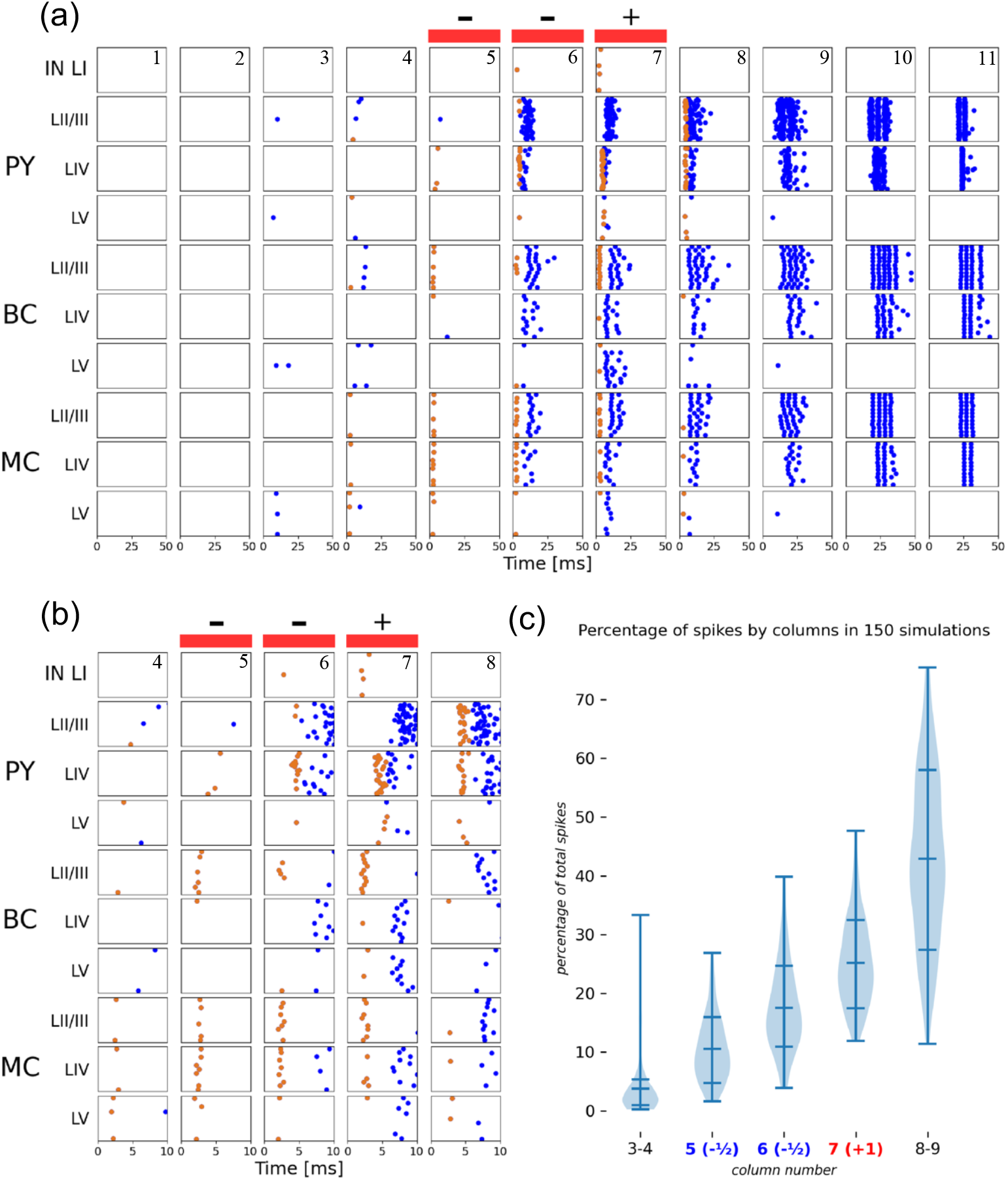
Directed propagation of pyramidal activity in raster plot of microcircuit trial. A raster plot displaying network behavior during and after stimulation in one trial of the microcircuit simulation across all cortical columns. Each cell within the microcircuit has its own coordinate on the y axis, and each dot is an action potential. Orange dots indicate spikes that are directly triggered by electrical stimulation (occurs during first 5 ms), while blue dots indicate spikes triggered via synaptic input. Figure 5a displays the first 50 ms of the simulation (the network is silent beyond this period), and Figure 5b zooms in on the first 10 ms in the five central columns. In Figure 5c, the number of spikes (in all cell types) was summed across the simulated cortical columns and averaged from 150 different simulations. In the vast majority of simulation trials, pyramidal activity in superficial layers propagates in a traveling wave to the right but is halted underneath the anodal electrodes by primarily Martinotti cell activity.

**Figure 6.**
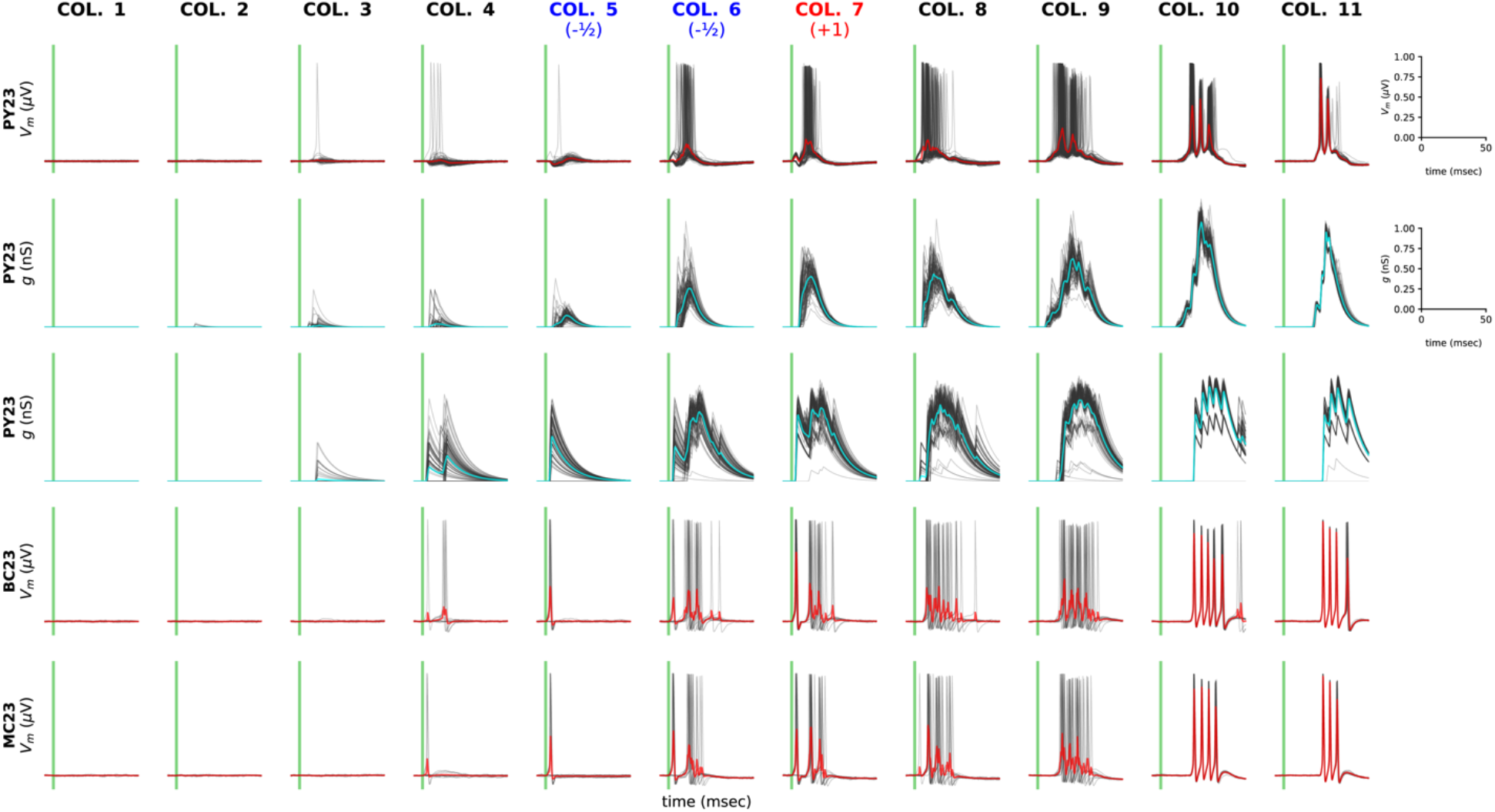
Voltage traces for layer II/III cells and PY input conductances during simulation show how inhibition causes directionality of traveling wave. Each of the subplots in the grid contain data for the collection of layer II/III cells specified by type and column. The x-axis for each subplot is time in milliseconds and is restricted to the interval from 5 msec before stimulation to 50 msec after. The stimulation period is depicted by the region in light green. Subplot columns, labeled at top, indicate the model columns; columns of 4 through 10 of the total 11 are included. The first three rows depict data from pyramidal cells, with the first row depicting the voltage trace, the second the total excitatory conductance (the sum of all incoming pyramidal connections), and the third the total inhibitory conductance (sum of inhibitory connections from layer I interneurons, basket cells, and Martinotti cells). The fourth and fifth rows depict the voltage traces of basket and Martinotti cells, respectively. In each subplot, there are light grey traces depicting each of the individual cells in the group, and their average (within the same column, layer, and subtype) in red. Sharp increases in the light grey trace indicate action potentials (spikes) and darker grey regions correspond to times when large numbers of cells spiked.

### Directionality of the stimulation-triggered wave can be explained by network inhibition

The activation probability curves in Figure 3 provide intuition into the network behavior during and immediately after the stimulation period (0-5 ms; Fig 5b). Let us first examine the column underneath the cathodal electrode (Fig 5b – column 7). While both layer II/III and layer IV pyramidal cells were predicted to be highly activated underneath the cathodal electrode (Fig. 3), only layer IV pyramidal cells were directly activated. Although all pyramidal cells were inhibited by moderate Martinotti activity, layer II/III pyramidal cells in particular were locally inhibited by strong synchronous LII/III basket cell activity because basket cell response dropped off with increasing cortical depth and because basket cells only inhibited within their own layer. Following stimulation, layer IV pyramidal cells excited layer II/III pyramidal cells and triggered a cluster of LII/III activity. There was negligible layer V excitation during stimulation and none following due to their low excitation probabilities and relatively small neuronal population.

Network behavior in response to anodal stimulation contrasted sharply with cathodal stimulation response and underpins unidirectional excitatory propagation (Fig 5b – columns 5 and 6). Although Layers II/III and IV pyramidal cells were still moderately activated by anodal stimulation (Fig 3a), very few cells were pushed above threshold due to strong inhibition. Martinotti cells showed a preference for anodal stimulation (according to Figure 3) and thus fired early in the stimulation period, inhibiting pyramidal cells across all cortical depths (since Martinotti cells universally synapsed to all pyramidal cells). This strong inhibitory force coupled with moderate superficial basket cell activity silenced almost all pyramidal activity across cortical layers.

In the column directly to the left of the electrode array (Fig 5b – column 4), there was negligible activity across all cell types and cortical layers. Not only did the electric field potential drop off significantly at this distance, but pyramidal and basket cells were already minimally activated by the anodal electrodes, and Martinotti cells were only moderately activated by the anodal electrodes due to their lack of myelination. In the absence of stimulating electric field potential or activating input from neighboring cortical columns, the first three columns exhibited no spiking activity at all (Fig 5b – columns 1-3). Hence, the excitatory pyramidal activity present underneath the electrodes did not propagate leftward past the anodal electrodes. This activity, however, did travel rightward past the electrode array, growing stronger and more synchronous as it propagated.

On the other side of the array, in the cortical column directly to the right of the cathodal electrode (Fig 5a & 5b – column 8), there was moderate direct activation of pyramidal cells and little direct activation of inhibitory cells. This follows from Figure 3, which depicts pyramidal cells continuing to be activated by the cathodal electrode. While basket cells were also moderately activated by cathodal stimulation, their joint inhibition with Martinotti cells was not enough to counter pyramidal stimulation response. This allowed for dense clusters of excitatory activity in pyramidal cells following stimulation.

In the second column to the right of the electrode array (Fig 5a – column 9), we see a dense cluster of highly synchronized pyramidal layer II/III activity that was slightly delayed from the activity in the column to its left. Although pyramidal cells were no longer directly stimulated in this cortical column, this activity resulted from the rightward cross-column propagation of excitatory signaling. Remarkably, this substantial, synchronized pyramidal activity grew more and more synchronous as the wave propagated rightward across cortical columns. This unidirectional propagation of excitatory signaling is an exceptional result of asymmetrical stimulation (Fig 5a – columns 9-11).

Following the stimulation period and initial clusters of activity that die down around 10 ms post-stimulation, there were a handful of waves of activity that ping-ponged between excitatory and inhibitory cells in columns with pyramidal excitation (Fig 5a – columns 6-11). In these columns, pyramidal activity activated both Martinotti and basket cells, which in turn inhibited pyramidal activity. This negative feedback loop went through a handful iterations over the course of a few milliseconds before halting pyramidal spiking entirely.

### Analysis of synaptic currents reveal mechanism of asymmetrical spiking activity

Next we analyzed voltage traces of individual neurons and synaptic dynamics to explain the causes of pyramidal asymmetrical spiking activity (Fig 6).

Voltage traces revealed that LII/III pyramidal cells were initially depolarized underneath the cathodal electrode (column 7), as they were highly activated by cathodal stimulation. However, a large inhibitory conductance immediately following the stimulation period dampened any stimulation-induced depolarization and hyperpolarized these pyramidal cells. The inhibitory conductance then fell and the excitatory conductance rose as layer II/III pyramidal cells gained excitatory input from neighboring layers and columns, bringing layer II/III pyramidal cells to threshold and triggering action potentials. The peak in inhibitory conductance midway through the pyramidal action potential was caused by pyramidal cell input into the inhibitory cells, which initiated a negative feedback loop that quickly subsided as pyramidal cells were silenced by inhibition. This effect occurred in all columns with substantial pyramidal activity.

Although there were similar inhibitory and excitatory conductance dynamics in the column underneath the central anodal electrode (column 6), the excitatory conductance was of a smaller magnitude overall and there were fewer action potentials because pyramidal cells were less excited by anodal stimulation. Underneath the leftmost electrode (column 5), moderate inhibitory conductance outweighed negligible excitatory conductance, leading to minimal pyramidal activity. There was little excitation in column 4 or any of the other leftward columns (not shown). On the other side of the electrode array, in column 8, high initial excitatory conductance and minimal inhibitory conductance resulted in strong initial pyramidal spiking. Pyramidal action potentials became more and more synchronous as they travel rightward, as evidenced by increasingly overlapped voltage traces. Together, these observations revealed the mechanisms of activity propagation to the right but not to the left in given stimulation settings.

## DISCUSSION

In this work we predict that an asymmetrical cortical stimulation protocol using a combination of anodal and cathodal electrodes may trigger propagating excitatory activity that shows strong directional preference. Our model has two steps: we first construct a biophysical model to predict activation probabilities across cell types in response to an asymmetrically applied electric field potential, and then incorporate these probabilities into a cortical microcircuit to model the network effects of stimulation. We found that pyramidal cells and basket cells are highly activated by the cathodal electrode and minimally activated by the anodal electrodes due to their myelination and horizontal axonal arbors, while Martinotti cells and Layer I interneurons are moderately activated by both electrodes but exhibit a slight preference for the anodal stimulation as they are unmyelinated and have vertically-oriented axonal arbors. Network model simulations revealed that this asymmetrical activation results in a traveling wave in superficial excitatory cells that propagates away from the electrode array, past the cathodal electrode, and into adjacent cortical columns, but does not propagate in the opposite direction past the leftmost anodal electrodes (Fig. 7).

**Figure 7.**
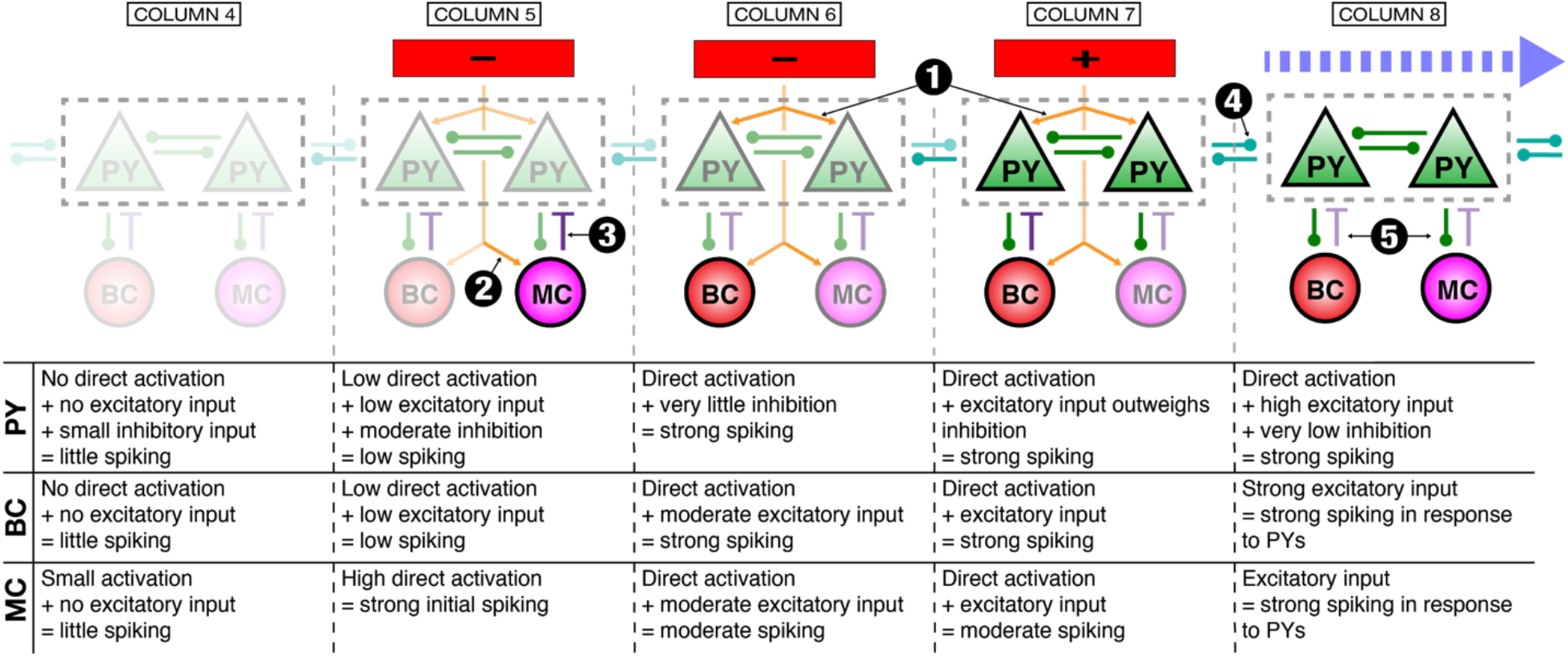
Summary of interactions resulting in unidirectional propagation. Key interactions are depicted graphically above and summarized in text in the table below. Columns 4 through 8 of the total 11 are included, indicated by labels at top. Cell and synaptic labels are as in Figure 4. Orange arrows coming from the electrodes and pointing toward cells indicate direct electrical stimulation, as opposed to synaptic inputs, depicted by the green, purple, and teal lines. The opacity of the cell labels approximately depicts the degree of spiking activity, with more transparent cells showing little or no spiking activity, and more opaque ones corresponding to more actively spiking cells. The opacity of lines indicates the strength of the input to cells, either synaptic or from the stimulating electrodes, e.g. more opaque orange arrows indicate that the cell being pointed to was more strongly stimulated directly. The light blue dashed arrow at top right indicates the initiation of synchronous, unidirectional propagation of activity in the direct right of the electrodes. Key events in initiation of this unidirectional propagation are indicated by the white numbers in black circles: ➀ Direct activation under (+) [col. 7] and central (-) [col. 6] induces strong spiking in PYs, BCs, MCs. ➁ Under left (-) [col. 5] direct activation induces strong spiking in MCs, but little in PYs, BCs. ➂ PYs in col. 5 further inhibited by MCs leads to little activity to left of electrode. ➃ Strong PY activity in cols. 6 and 7 propagates rightward via cross-column PY-PY synapses, overcoming moderate inhibition. ➄ PY activity causes spiking in MCs and BCs as it propagates rightward, feedback between PYs and inhibitory cells causes increasingly synchronous spiking. Layer I interneurons have been omitted, as they did not contribute strongly to the propagation described here.

This model develops a stimulation protocol that can be experimentally verified in future studies. We found that inhibitory cells are the cause of unidirectional propagation, which implies that excitatory activity would exhibit unhindered propagation without inhibition. Inhibitory cells could be deactivated optogenetically during stimulation to test model predictions. Martinotti cells may also be deactivated separately from basket cells via genetically-targeted optogenetics to localize the main source of inhibition.

### Traveling waves in the brain

Experimental recordings have observed traveling waves during most types of cortical processing. They relay information across a range of distances and thereby coordinate processes such as memory, perception, language, orientation, executive functions, and more across distant brain regions^17,27–29^. Multi-electrode recordings in human and animal subjects have demonstrated the ubiquity of traveling waves in cognitive function^17^. A growing body of evidence therefore places traveling waves as integral to cognition, as they facilitate fast, efficient information transfer and local-to-global communication^30^. Multi-electrode recordings show that propagating activity present in the human cortex is often directional, traveling from one point to another. The ability to generate directional propagation via stimulation would allow for unique precision in and control over induced activity. This has widespread implications for the restoration or enhancement of high-level brain function, particularly because many neuropsychiatric disorders are marked by abnormal or absent propagating activity.

In the model, propagating activity was confined to supragranular cortical layers. This may not be a limitation if the goal is to reproduce natural waves, inasmuch as spontaneous traveling waves in the human cortex have also been found to be largely confined to upper layers, including the alpha rhythm during waking^18^, and spindles and slow oscillation during sleep^31,32^.

Traveling waves have long been studied with a variety of computational models, and numerous mechanisms have been proposed to explain how activity propagates across neuronal networks^33,34^. Cortical propagating waves that are triggered specifically by electrical stimulation have been recorded in mammalian cortical slices^35–37^, as well as in non-human mammals^38–40^ and simulated in non-mammalian computational models^41^, but remain understudied in human subjects. This previous experimental work, both *in vivo* and *in silico*, has yielded scarce evidence of asymmetrical travelling wave propagation or reliable wave generation analogous to that reported here. To the extent that previous work has focused on travelling wave propagation initiated by stimulation^42^, these studies examined stimulation through the skull and meninges, and are thus not directly comparable to this model of intracranial stimulation of the cortical surface. Many computational models exist that model the effects of stimulation on the brain, including some that have constructed Hodgkin-Huxley microcircuits^43,44^ and modeled cortical surface stimulation^45^. However, few have modeled multi-electrode or asymmetrical stimulation or reproduced traveling waves using surface electrodes. Many existing stimulation models have focused on stimulation of a particular nerve^46,47^ or isolated cell type^48,49^, as opposed to the functioning cortical microcircuit presented here, which can be more readily adapted to other cortical regions. In addition, stimulation has more often been simulated in these models by a simple application of superthreshold current or a uniform electrical field^50,51^. Thus, by combining the two-phase biophysical model with asymmetrical, multi-polar surface stimulation, our approach synthesizes existing achievements into a single coherent, clinically-adaptable model that uniquely sheds light on the generation of propagating wave activity.

### Clinical relevance of our findings

Brain stimulation is becoming increasingly common in clinical and experimental settings, especially using multi-electrode arrays^52^. As such, it is pressing that we develop accurate models of the effects that multi-electrode stimulation has on neural activity. While sometimes the explicit goal of stimulation may be to disrupt aberrant activity to restore normal functioning, increasingly the goal is to induce the desired brain activity directly via stimulation, as our work demonstrates.

Changes in neural plasticity result from patterned activity, with the particular changes in connectivity contingent on the specific timing and order of activity^53^. Stimulation protocols that induce neural activity which continues past the stimulus duration are more likely to alter cellular and synaptic properties in favor of the induced activity, in contrast to stimulation protocols that briefly activate broad swaths of cells without triggering existing activity patterns. Thus, initiating propagating waves within tailored spatiotemporal constraints is a promising way to retrain neural networks and enhance or silence brain functions in a targeted way.

The generation of traveling waves may serve as a promising therapy for a variety of neurological disorders. For example, it has been previously suggested that triggering propagating activity in perilesional areas where waves are otherwise aberrant or absent may be an effective therapy for post-stroke aphasia^54^. While the ultimate clinical applications of this technique are uncertain, stimulation-induced traveling waves may have the potential to offset inhibition in cortical spreading depression^55,56^, reduce the risk of seizure while determining which brain tissue to remove from epileptic patients^11^, and enhance memory formation when applied during learning or recall periods^13,14,16,57,58^, as consistency in traveling wave direction is positively correlated with working memory efficiency^59^.

### Limitations

In this work, we modelled a single, short stimulating pulse. However, clinical stimulation is most often composed of longer pulses or pulse trains and is usually performed with bipolar electrodes delivering biphasic pulses in order to prevent damaging Faradic currents^60,61^. These stimulation paradigms modulate properties over time that are not accounted for in the current model such as underlying dendritic and axonal dynamics as well as synaptic interactions. Thus, our biophysical approach may be expanded in future studies to incorporate these steady-state properties through alternative modeling approaches such as the cylinder model^62,63^ or the multi-compartmental model^64^. However, we chose to model the activation probability of the axonal instead of the dendritic arbor in this work because experimental evidence shows that the nodes of Ranvier, followed by the axon hillock, are the most excitable neuronal elements by far via direct stimulation^65–68^ as they both have a high concentration of sodium channels^69^. In contrast, direct stimulation of the dendritic arbor generates transmembrane currents that propagate to the axon hillock, but these effects are strongly attenuated and delayed, and are negligible compared to direct stimulation of the nodes of Ranvier and axon hillock.

In the microcircuit phase of the model, the connectivity between different cell types follows a canonical microcircuit model. While this approach characterizes the main signal pathways and feedback loops present within cortical columns^70–72^, finer details are not modeled such as the variability of layer II/III pyramidal cells and their projections^73,74^, descending projections to inhibitory cells from excitatory cells^75^, or the contribution of less common interneuron cell types. These and other characteristics such as axonal and dendritic arborization change between species and cortical areas. The cells within the cortical microcircuit model could be extended from single-to multi-compartment neurons^76^ in order to distinguish tuft versus soma-targeting interneurons, which may further differentiate the inhibitory power of interneuron cell types^24^. This phase of the model may be further expanded from a two-dimensional plane to a three-dimensional circuit in the volume of cortex underneath the electrodes, which would allow for a better understanding of how activity spreads across space.

### Generalization of this approach

While this work modeled a specific stimulation protocol, the approach is generalizable to a variety of clinical applications. The model can be customized in terms of the number and shape of the electrodes, the arrangement of anodal and cathodal electrodes, the intensity of the applied current, and the brain area studied (provided sufficient neuronal reconstructions are available). This model can also be modified to simulate other applied stimulating pulses as long as an activation threshold has been experimentally determined. However, further adjustments are necessary to adapt this model to human stimulation protocols such as modeling a biphasic bipolar pulse, incorporating human cell reconstructions as they become available, and adjusting model parameters to account for the thicker human pia and cortex.

### Conclusions

This work models an asymmetrical stimulation paradigm that could be implemented to initiate unidirectional traveling waves in the cortex. A biophysical model is integrated with a network-computational model to predict the behavior of single neurons as well as the cortical network dynamics resulted from multielectrode stimulation. This model provides hypotheses and stimulation paradigms that can be verified experimentally and also provides an avenue for further experimental work to be guided systematically rather than in blind trial-and-error fashion. These results demonstrate how complex stimulation protocols could be harnessed to generate persistent changes in activity with the potential to restore normal brain function in neurological and psychiatric conditions.

## Methods

### Cell reconstruction selection

All neuronal cell reconstructions were chosen from publicly available datasets on neuromorpho.org^22,25,77–80^. The types of cells included and their respective datasets are listed in table S1. We used multiple cell reconstructions for each cell type to account for anatomical diversity. We used experimental measurements to approximate the cutoff depths for each layer (Fig 1a)^24,25^.

### Calculating the electric field potential generated by the electrode array

The electrode array modeled in this study is composed of three square electrodes (each 150 µm by 150 µm) placed linearly on the surface of the cortex. Two electrodes have negative current (- 75 µA each) and one electrode has positive current (150 µA) (Fig 1a), and stimulation is applied for 200 μs. These values are in accordance with common experimental parameters^21^. Assuming that the current sources are homogenous square electrodes, we calculated the electric field potential of each electrode as follows:

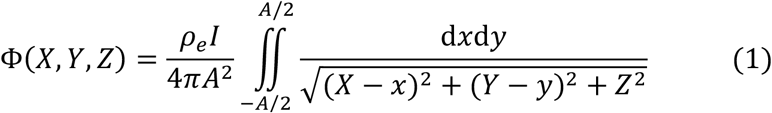

*I* is net current, *ρ*_e_ is extracellular resistivity and A is the length of the square electrode edge (Fig.1a). In this study, A=150 μm and net current I is either −75 μA or 150 μA. The derivation of this formula can be found in Komarov et al.^20^.

We sum all three electric field potentials at each point in space to determine the overall electric field potential.

### Estimating the activating function

Derived from one-dimensional cable theory^63,66^, the transmembrane voltage dynamics of axonal segments can be modeled as follows:

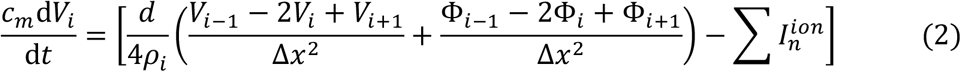

*V*_*i− 1*_, *V*_*i*_, and *V*_*i+1*_ are transmembrane voltages of the neighboring axonal compartments (sub index indicates number of the compartment), *c*_*m*_ *=* 1 μF/cm^2^ is a capacitance of membrane per square unit area, d is the diameter of the axon (typically between 10 μm and 1 μm), *ρ*_*i*_ = 300 Ω·cm is a resistivity of axoplasm, and Δ*x* is a discretization parameter that defines length of the compartment. The term 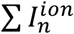 is the sum of intrinsic ionic currents such as leak currents and fast potassium and sodium currents for spike generation. From above, the activating function is the effective transmembrane current that arises due to extracellular electrical stimulation and is described as

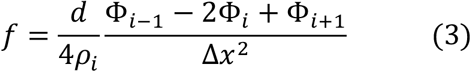

where Φ_*i− 1*_, Φ_*i*_, and Φ_*i+1*_ stand for extracellular potentials in the vicinity of axonal compartments. In the limit Δ*x* → 0 the expression becomes 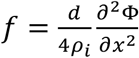, where the *x* axis represents the direction of the axonal fiber. To estimate the probability of axonal activation, we calculate the transmembrane current generated in each small component piecewise along reconstructed axons using the activating function. This calculation is based upon the orientation of each component in space because the activating function is computed along the axonal direction by definition. Since jitters may be present along the edges of reconstructions, we estimate component direction using neighboring segments in space up to 10 microns away.

### Estimating the threshold of activation

After calculating the transmembrane current generated within each axonal segment of cell reconstructions, we used a threshold of activation to determine whether a neuronal response was triggered. This threshold was drawn from *in vivo* experiments which define the threshold injected current *I* required to induce a threshold effective current *f* at the axonal initial segment a distance *d* from the electrode^48,81^. Komarov et al.^20^ simulated one such experiment by computing the activation current *f* at the axon initial segment of a layer II/III pyramidal cell while varying the stimulation current *I* and distance *d*. This simulation used a 200 μs duration stimulating pulse, which is typical of similar *in vivo* experiments^48,81^. The value *f = f*_*th*_ = 3 pA/μm^2^ fully replicated the experimentally observed current-distance relationship across varying stimulation currents and distances. Thus, this threshold value *f*_*th*_ was used to determine whether each axonal segment was activated by induced transmembrane current.

### Computing the average activation probability per cell type

To determine whether a given neuronal reconstruction would be activated by applied current, we used the activating function to calculate how many axonal segments had above-threshold transmembrane current values that could initiate an axonal action potential (assumed at nodes of Ranvier); these above-threshold axonal segments are collectively called the trigger area. The activating function threshold was set to *f*_*th*_ = 3 pA/μm^2^ for myelinated axons but to *f*_*th*_ = 60 pA/μm^2^ for unmyelinated axons, since unmyelinated segments are significantly less excitable as they have fewer sodium channels^61^. In this model, we assumed pyramidal and basket cells are myelinated and Martinotti and Layer I interneurons are unmyelinated based upon experimental data^25,75,82^. For each cell reconstruction at a given point in space (i.e. a given coordinate in the x-z plane, Fig 1a), we found the probability of firing as follows:

We computed the activating function along the axonal segments of the cell reconstruction and found the trigger area *L* as described above.

a. For myelinated axons, we assumed that at least one node of Ranvier must be present in the trigger area in order to initiate an axonal response. To approximate the probability of a node of Ranvier in the trigger area, we first discretized the trigger area into segments that are the typical length of a node of Ranvier (*k*=1 μm). If we approximate the mean internodal distance as D = 100 μm, then *p*_*n*_ *= k/D* is the probability that there is a node of Ranvier in a given segment. It follows that (1 – *p*_*n*_)^*N*^ is the probability that the trigger area *L* does not contain a node of Ranvier, where *N* = *L*/ *N* is the number of segments of length *L* that can fit into a fiber of total length *L*. Therefore, the overall probability p that a myelinated axonal arbor will be activated is estimated as 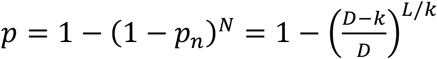.
b. For unmyelinated axons, we assumed that any axonal segment with an activating function above threshold (*L* > 0) resulted in an overall cell response, since the entire membrane is exposed to extracellular space.

Given that variation in cell position and orientation within each cortical layer is present in nature, we additionally average across neuronal rotations and vertical shifts. Thus, we computed the average activation probability for each cell type/cortical layer as follows: for each cell reconstruction, we first position the soma at a point along the x-axis within its cortical layer. We then performed four rotations about the vertical axis of the cell reconstruction in three-dimensional space, and at every rotation we calculate the likelihood of activation by computing the activation function across the axonal arbor, as outlined above. We then averaged across these four probabilities and set the result as the activation probability for said point. After this, we incrementally shift the soma of the reconstruction vertically within its cortical layer and find the mean activation probability at each point. We averaged across all vertical shifts to obtain an approximate spiking probability for this cell reconstruction at this point along the x-axis. We then computed the activation probability of all cell reconstructions across the x-axis in this manner, and finally average across all reconstructions within a given cell type and cortical layer to find the expected spiking probability in response to stimulation for each class of cell.

### Computational model of the cortical circuit

#### Rationale

The network model is composed of eleven interconnected cortical columns, and each column contains layer I interneurons and pyramidal, basket, and Martinotti cells from layers II-V (same as the biophysical analysis). The number of cells within each column is outlined in table S2. This balance of excitatory to inhibitory cell types approximates the true cell composition of the rat cortex, where pyramidal cells are the primary excitatory cells and basket and Martinotti cells comprise the majority of inhibition within and across layers^24,25^. Cells were constructed to only spike if receiving synaptic input or electrical stimulation. Each cell behaves according to Hodgkin-Huxley dynamics, with a handful of parameters differentiating excitatory and inhibitory cells. Basket cells were modeled as fast-spiking cells, while the other cell types were modeled as regular-spiking cells with spike rate adaptation. Inhibitory cells were fired more quickly than excitatory cells in response to activation because all interneurons were modeled as having a lower leak current^56^. Some additional parameters are: a fast Na^+^-K^+^ spike generating mechanism (all cells), high-threshold activated Ca^2+^ current (for pyramidal cells), and slow calcium-dependent potassium (AHP) current (for regular spiking cells).

#### Equations

The membrane potential is described by following equation^83^:

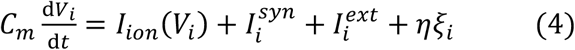

The ionic currents *I*_*ion*_(*V*_*i*_), which are responsible for intrinsic cells dynamics, are as follows:

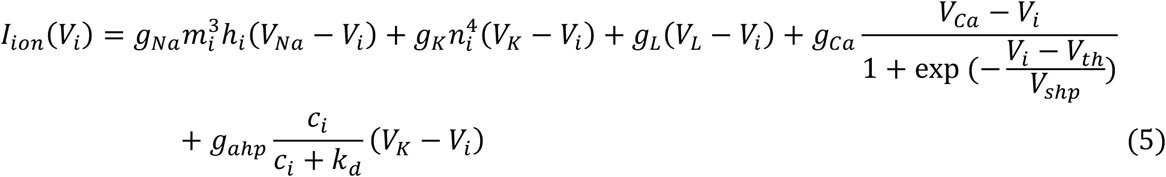

The gating variables *m*_*i*_, *n*_*i*_, *h*_*i*_ evolve according to:

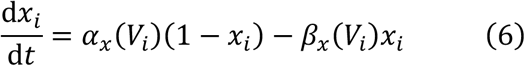

where *x*_*i*_ is one of gating variables. The functions *α*_x_(*V*) and *β*_x_(*V*) are:

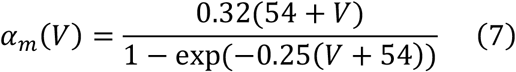

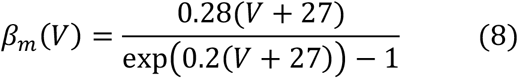

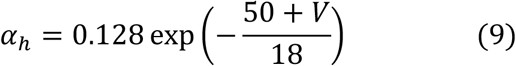

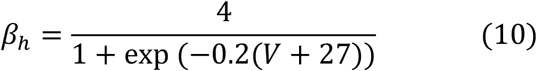

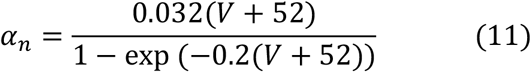

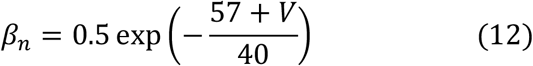

Calcium concentration *c*_*i*_ obeys the following equation:

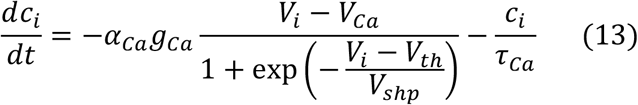

and governs calcium-dependent hyperpolarizing potassium current

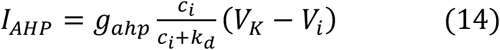

which is responsible for spike-frequency adaptation. The synaptic input was modeled according to:

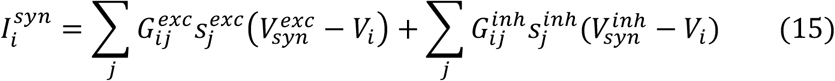

where synaptic variables 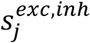 are governed by the following equation:

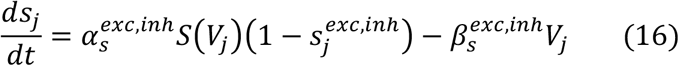

The function *S*(*V*) reads:

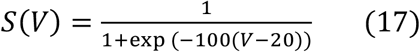

Initially, the network runs without stimulating input for 200 ms to simulate pre-existing activity. Then the network is stimulated and runs for an additional 500 ms. To simulate electrical stimulation, we used the binary term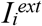to inject above-threshold current into a subset of randomly-chosen neurons within each cell type/cortical layer such that the fraction of neurons induced to spike corresponds to the activation probabilities calculated in the biophysical phase of the model. The term *ηξ*_*i*_(*t*) models spontaneous background activity as a white noise process (*ξ*) with standard deviation *η*. All model parameters are listed in Table S2 (unless specified in the description of simulations), and the network structure and connectivity are described in Table S3 and Table S4 respectively. Cells were synaptically coupled by excitatory (AMPA) and inhibitory (GABA_A_) connections. The strength and probability of connections between layers and cell types are set according to a canonical cortical circuit^75^.

## Acknowledgements

This work was supported by NIH-NINDS (grant 1R01NS19553).

## SUPPLEMENTARY TABLES

**Table S1:** Summary of datasets with reconstructed cells.

| Cell type | # of Reconstructions | Reference | Strain (Age) |
| --- | --- | --- | --- |
| Pyramidal cells (Layer II/III) | 21 | [74] | Wistar (P20-25) |
| Pyramidal cells (Layer IV) | 11 | [78] | Wistar (P19-21) |
| Pyramidal cells (Layer Va) | 14 | [79] | Wistar (P20-21) |
| Basket cells (Layer II/III) | 96 | [81] | Wistar (P13-15) |
| Basket cells (Layer IV) | 82 | [81] | Wistar (P13-15) |
| Basket cells (Layer V) | 57 | [81] | Wistar (P13-15) |
| Martinotti cells (Layer II/III) | 13 | [23] | Wistar (P13-16) |
| Martinotti cells (Layer V) | 7 | [23] | Wistar (P13-16) |
| Horizontal interneurons (LI) | 59 | [80] | Wistar (P13-16) |
| Descending interneurons (LI) | 29 | [80] | Wistar (P13-16) |
| Small interneurons (LI) | 27 | [80] | Wistar (P13-16) |

**Table S2:** Model parameters for network simulations.

| Parameter | Description | Value |  |  |
| --- | --- | --- | --- | --- |
| $C_m$ ($\mu\text{F}/\text{cm}^2$) | Membrane capacitance | 1 | | |
| $\eta$ | Standard deviation of spontaneous background activity | 0.4 | | |
| $V_{Na}$ (mV) | $\text{Na}^+$ reversal potential | 50 | | |
| $V_K$ (mV) | $\text{K}^+$ reversal potential | -90 | | |
| $V_{Ca}$ (mV) | $\text{Ca}^{2+}$ reversal potential | 120 | | |
| $V_{shp}$ (mV) | Slope of $\text{Ca}^{2+}$ activation function | 2.5 | | |
| $V_{th}$ (mV) | Threshold potential | -25 | | |
| $k_d$ | Equilibrium dissociation constant | 1 | | |
| $\alpha_{Ca}$ ($\text{ms}^{-1}$) | $\text{Ca}^{2+}$ activation time constant | 0.005 | | |
| $\tau_{Ca}$ ($\text{ms}^{-1}$) | $\text{Ca}^{2+}$ decay time constant | 200 | | |
| $\alpha_s^{exc}$ ($\text{ms}^{-1}$) | Excitatory synaptic activation time constant | 0.6 | | |
| $\alpha_s^{inh}$ ($\text{ms}^{-1}$) | Inhibitory synaptic activation time constant | 0.5 | | |
| $\beta_s^{exc}$ ($\text{ms}^{-1}$) | Excitatory synaptic deactivation time constant | 0.2 | | |
| $\beta_s^{inh}$ ($\text{ms}^{-1}$) | Inhibitory synaptic deactivation time constant | 0.1 | | |
| $V_{syn}^{exc}$ (mV) | Excitatory synaptic reversal potential | 0 | | |
| $V_{syn}^{inh}$ (mV) | Inhibitory synaptic reversal potential | -80 | | |
| Cell-type specific parameters |  | PYs | BCs | MCs |
| $g_{Na}$ ($\text{mS}/\text{cm}^2$) | $\text{Na}^+$ channel conductance | 30 | 60 | 60 |
| $g_K$ ($\text{mS}/\text{cm}^2$) | $\text{K}^+$ channel conductance | 2.5 | 10 | 10 |
| $g_L$ ($\text{mS}/\text{cm}^2$) | Leak channel conductance | 0.6 | 0.6 | 0.4 |
| $g_{Ca}$ ($\text{mS}/\text{cm}^2$) | $\text{Ca}^{2+}$ channel conductance | 0.1 | 0 | 0.1 |
| $g_{ahp}$ ($\text{mS}/\text{cm}^2$) | Slow $\text{Ca}^{2+}$-dependent (after-hyperpolarization) $\text{K}^+$ channel conductance | 0.5 | 0 | 0.75 |
| $V_L$ (mV) | Leak channel reversal potential | -67 | -65 | -65 |

**Table S3:** Structure of the network.

| Cortical cell type | Layer | Cells/column | Total cells |
| --- | --- | --- | --- |
| IN | I | 12 | 132 |
| PY | II/III | 100 | 1100 |
| BC | II/III | 100 | 1100 |
| MC | II/III | 24 | 264 |
| PY | IV | 12 | 132 |
| BC | IV | 12 | 132 |
| MC | IV | 12 | 132 |
| PY | V | 12 | 132 |
| BC | V | 12 | 132 |
| MC | V | 12 | 132 |
| *total* |  | 308 | 3388 |

**Table S4:** Connectivity within the network.

| Presynaptic |  | Postsynaptic |  | Cross-column | Type | Strength | Probability |
| --- | --- | --- | --- | --- | --- | --- | --- |
| Type | Layer | Type | Layer |  |  |  |  |
| PY | II/III | PY | II/III | TRUE | AMPA | 0.4 | 0.1 |
| PY | IV | PY | IV | TRUE | AMPA | 0.4 | 0.1 |
| PY | V | PY | V | TRUE | AMPA | 0.4 | 0.1 |
| PY | II/III | PY | II/III | FALSE | AMPA | 0.75 | 0.1 |
| PY | II/III | BC | II/III | FALSE | AMPA | 0.75 | 0.1 |
| PY | II/III | MC | II/III | FALSE | AMPA | 0.75 | 0.1 |
| PY | II/III | PY | IV | FALSE | AMPA | 0.25 | 0.05 |
| PY | IV | PY | II/III | FALSE | AMPA | 1.5 | 0.05 |
| PY | IV | PY | IV | FALSE | AMPA | 0.75 | 0.1 |
| PY | IV | BC | IV | FALSE | AMPA | 0.75 | 0.1 |
| PY | IV | MC | IV | FALSE | AMPA | 0.75 | 0.1 |
| PY | IV | BC | II/III | FALSE | AMPA | 1.5 | 0.05 |
| PY | IV | MC | II/III | FALSE | AMPA | 1.5 | 0.05 |
| PY | V | PY | V | FALSE | AMPA | 0.75 | 0.1 |
| PY | V | PY | II/III | FALSE | AMPA | 0.5 | 0.05 |
| PY | V | BC | V | FALSE | AMPA | 0.75 | 0.1 |
| PY | V | MC | V | FALSE | AMPA | 0.75 | 0.1 |
| MC | II/III | PY | II/III | FALSE | GABA | 0.75 | 0.05 |
| MC | II/III | PY | IV | FALSE | GABA | 0.75 | 0.05 |
| MC | II/III | PY | V | FALSE | GABA | 0.75 | 0.05 |
| MC | IV | PY | II/III | FALSE | GABA | 0.75 | 0.05 |
| MC | IV | PY | IV | FALSE | GABA | 0.75 | 0.05 |
| MC | IV | PY | V | FALSE | GABA | 0.75 | 0.05 |
| MC | V | PY | II/III | FALSE | GABA | 0.75 | 0.05 |
| MC | V | PY | IV | FALSE | GABA | 0.75 | 0.05 |
| MC | V | PY | V | FALSE | GABA | 0.75 | 0.05 |
| BC | II/III | PY | II/III | FALSE | GABA | 1.5 | 0.25 |
| BC | IV | PY | IV | FALSE | GABA | 1.5 | 0.25 |
| BC | V | PY | V | FALSE | GABA | 1.5 | 0.25 |
| MC | V | PY | IV | FALSE | GABA | 1 | 0.2 |
| MC | V | PY | V | FALSE | GABA | 1.5 | 0.25 |
| IN | I | PY | II/III | FALSE | GABA | 0.3 | 0.3 |
| IN | I | PY | IV | FALSE | GABA | 0.3 | 0.3 |
| IN | I | PY | V | FALSE | GABA | 0.3 | 0.3 |
| PY | II/III | IN | I | FALSE | AMPA | 0.3 | 0.3 |

